# Emergent intra-pair sex differences and behavioral coordination in pair bonded prairie voles

**DOI:** 10.1101/2021.09.03.458892

**Authors:** Liza E. Brusman, David S. W. Protter, Allison C. Fultz, Maya U. Paulson, Gabriel D. Chapel, Isaiah O. Elges, Ryan T. Cameron, Annaliese K. Beery, Zoe R. Donaldson

## Abstract

In pair bonding animals, coordinated behavior between partners is required for the pair to accomplish shared goals such as raising young. Despite this, experimental designs rarely assess the behavior of both partners within a bonded pair. Thus, we lack an understanding of the interdependent behavioral dynamics between partners that likely facilitate relationship success. To identify intra-pair behavioral correlates of pair bonding, we used socially monogamous prairie voles, a species in which females and males exhibit both overlapping and distinct pair bond behaviors. We tested both partners using social choice and non-choice tests at short- and long-term pairing timepoints. Females developed a preference for their partner more rapidly than males, with preference driven by different behaviors in each sex. Further, as bonds matured, intra-pair behavioral sex differences and coordinated behavior emerged – females consistently huddled more with their partner than males did, and partner huddle time became correlated between partners. When animals were allowed to freely interact with a partner or a novel in sequential free interaction tests, pairs spent more time interacting together than either animal did with a novel. Pair interaction was correlated with female, but not male, behavior. Via a social operant paradigm, we found that pair-bonded females, but not males, are more motivated to access and huddle with their partner than a novel vole. Together, our data indicate that as pair bonds mature, sex differences and coordinated behavior emerge, and that these intra-pair behavioral changes are likely organized and driven by the female animal.

## Introduction

Interpersonal relationships require social cooperation to achieve shared goals, such as in socially monogamous pair bonds where two individuals share resources and offspring care. Because of these shared responsibilities and the lack of ongoing mate selection, monogamous species are often thought to exhibit fewer sex differences (Kleiman 1977; Sue Carter *et al*. 1995). However, there are well documented examples of behavioral sex differences in monogamous species (DeVries *et al*. 1996; DeVries & Carter 1999; Bales & Carter 2003; Prounis *et al*. 2018) which, unlike those observed in non-monogamous species, may emerge after a pair bond has formed to facilitate intra-pair cooperation and ensure reproductive success.

Among monogamous prairie voles, there are sex differences in parental care (Gruder-Adams & Getz 1985; Lonstein & De Vries 1999). Females and males exhibit similar parental behaviors, but they display these behaviors to different degrees across pup development and across subsequent litters (Solomon 1993; Lonstein & De Vries 1999; Rogers *et al*. 2018). By “trading off” duties, prairie vole parents can provide more active care for their pups, which promotes the pups’ physiological and behavioral development (Bales *et al*. 2007; Ahern & Young 2009; Rogers *et al*. n.d.). However, whether *intra-pair* coordinated sex differences in partner-directed behavior emerge as a function of relationship formation and maturation remains unexamined, especially as the vast majority of studies focus on only one member of a pair.

In addition to biparental care, prairie vole pair bonds are hallmarked by an affiliative partner preference that develops more rapidly in females than in males (DeVries & Carter 1999). Here, we characterized the social behavior of both members of bonded pairs at short-term and long-term timepoints post-pairing. We employed complementary choice and non-choice social tests, showing that organization of intra-pair behavior emerges as a function of bond duration, with coordinated but distinct changes occurring in each sex.

While partner-affiliative behavior is the gold standard for determining whether a pair bond has formed, these tests do not separate the appetitive and consummatory aspects of partner and novel interaction. To deepen our understanding of the underlying behavioral mechanisms that drive sex differences in pair bond behavior, we tested partner- and novel-directed social motivation in pair bonded voles using an operant choice, fixed ratio lever-pressing task. In accordance with a prior report using a non-choice-based, progressive ratio operant task, we found that females exhibited greater partner-directed motivation than males (Beery *et al*. 2021). Together, this work has important implications for deepening our understanding of social behaviors by uncovering behavioral mechanisms that reinforce pair bonds and delineating the interdependent dynamics *between* partners that facilitate relationship success.

## Results

We tested both members of bonded pairs in the partner preference test (PPT) and the free interaction test, at short- and long-term pairing timepoints (Fig. 1A), enabling us to identify consistent intra-pair sex differences that emerge as a function of bond maturation and examine how pair bonds develop over time. Accordingly, all PPT and free interaction metrics were specifically analyzed in a pairwise fashion using repeated-measures ANOVA (RM-ANOVA). P-values reported below represent post-hoc paired t-tests with Bonferroni correction with all additional statistics available in Supplementary Table S1.

**Figure 1:**
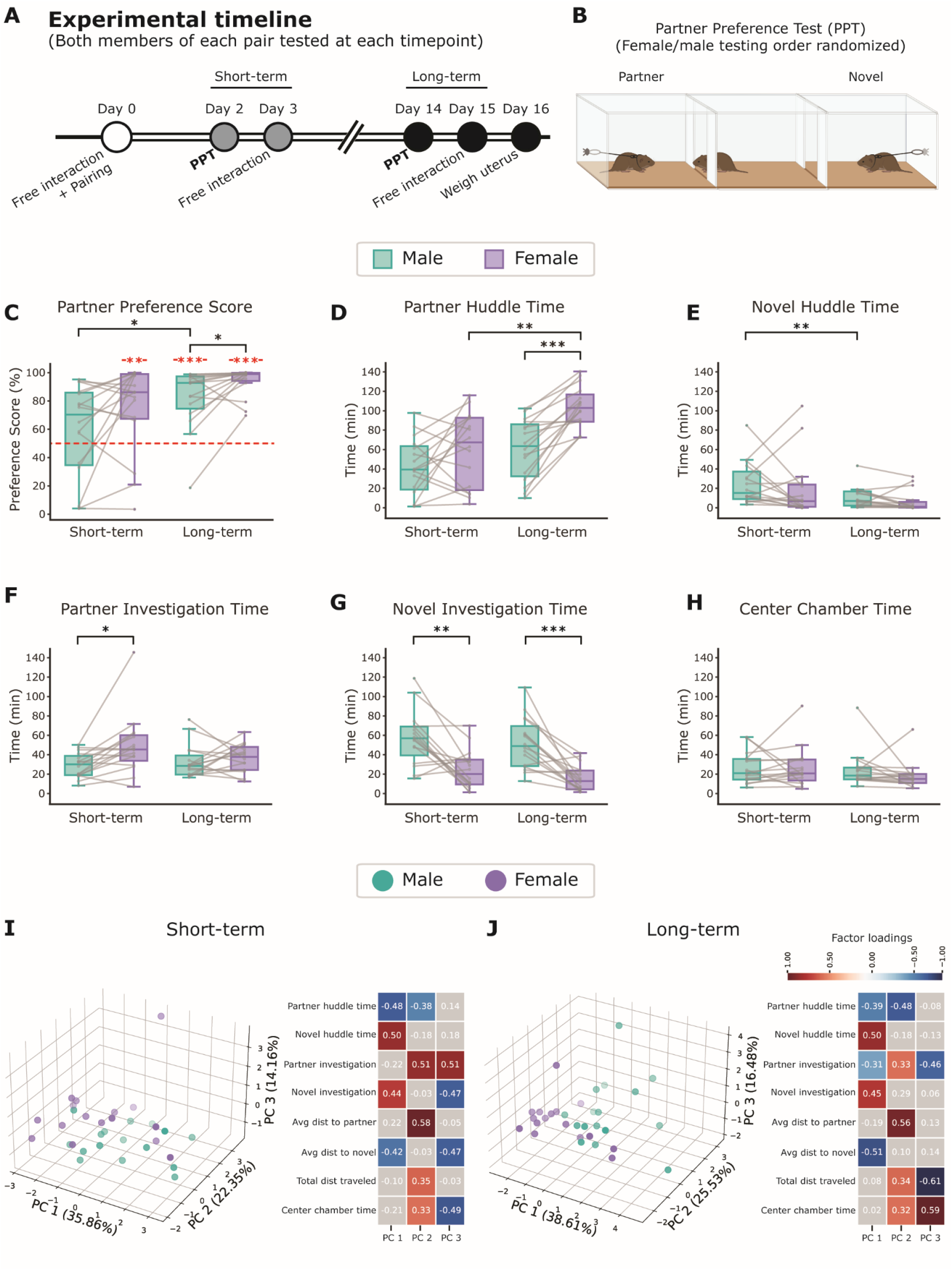
Sex differences in partner preference metrics. **A.** Schematic of experimental timeline. Animals (n = 16 F, 16 M) underwent a free interaction period with two novel animals: their eventual partner and a non-partner, before being paired for the remainder of the experiment. Partner preference tests (PPTs) were conducted 2 days (short-term) and 14 days (long-term) after pairing. **B.** Diagram of partner preference test. **C.** Partner preference scores for females and males at short and long term timepoints, calculated for each animal as Partner Huddle Time / (Partner Huddle Time + Novel Huddle Time) x 100%. Red asterisks denote significant difference from the null hypothesis of no partner preference (50%) using a one-sample t-test. Females form partner preferences by the short-term timepoint, while males do not (females: p = 0.0049, males: p = 0.28). By the long-term timepoint, both females and males display a partner preference (females: p = 3.3×10^-11^, males: p = 2.1×10^-5^) Males show an increase in partner preference between short- and long-term (p = 0.044). Sex differences are not apparent at the short-term timepoint (p = 0.30) but emerge by the long-term timepoint (p = 0.037). **D.** Total partner huddle duration for females and males at short- and long term timepoints. Females huddled more than their male partner at long (p = 3.2×10^-5^) but not short term timepoints (p = 0.45). Only females increase their partner huddle time between short- and long-term (p = 0.0052). **E.** Total novel huddle duration. No sex differences were observed at either timepoint, but males show decreased novel huddle time at the long-term compared to short-term timepoint (p = 0.0064). **F.** Partner investigation time, calculated as Partner Chamber Time minus Partner Huddle Time. Females investigated their partner more than males, but only at the short-term timepoint (short-term: p = 0.028, long term: p = 1.00). **G.** Novel investigation time. Males spent more time investigating the novel than females at both timepoints (short term: p = 0.0047, long term: p = 6.8×10^-4^). **H.** Time in the center chamber. No sex differences of time-dependent changes were observed for time in the center chamber (short term: p = 1.00, long term: p = 0.58). **I.** Principal component analysis (PCA) and factor extraction of mutually exclusive partner preference metrics at short term timepoint. Females and males are largely overlapping in the PCA. **J.** PCA and factor extraction of partner preference metrics at long term timepoint. Females cluster separately from males.

### Consistent intra-pair sex differences in PPT behavior emerge as a function of bond maturation

We first examined social behavior metrics in the classic partner preference test in both members of pairs following short-term (2 days) and long-term (2 weeks) cohabitation (Fig 1A). Traditionally, the ratio of partner to novel huddle time has been used to infer pair bond status. We calculated a partner preference score (Partner Huddle / (Partner + Novel Huddle)) to determine whether pair bonds had formed. Compared to a null value of 50%, we found that at the short-term timepoint, only females display a partner preference (females: p = 0.0049, males: p = 0.28), but both females and males have a partner preference at the long-term timepoint (Fig 1C, females: p = 3.3×10^-11^, males: p = 2.1×10^-5^). Further, there is an increase in partner preference score between the short- and long-term timepoints for males (p = 0.044), but not females (p = 0.42). This is consistent with prior data indicating that males take longer than females to establish a partner preference (DeVries & Carter 1999).

We next asked what specific behaviors within the PPT contribute to our observed sex difference in the emergence of partner preference. When looking at raw huddle time, we saw that females, but not males, increase their partner huddle time as they transition from short-to long-term timepoints (Fig 1D, females: p=0.0052, males: p = 0.37). Conversely, males huddled less with the novel at the long-term timepoint compared to the short-term timepoint, while females huddled the same amount with the novel at both timepoints (Fig 1E, females: p = 0.60, males: p = 0.0064). Accordingly, we can conclude that the formation and strengthening of partner preference over time occurs via different behavioral processes in females and males. Females increase partner-directed huddle even after a partner preference has already developed, while males decrease novel-directed huddle without a significant change in partner-directed affiliation, driving an increase in partner preference. The latter replicates prior work (Harbert *et al*. 2020).

We followed our longitudinal analysis of male and female behavior by asking whether there were intra-pair sex differences within either timepoint. There was no difference in partner preference score between females and males at the short-term timepoint (p = 0.30), but by the long-term timepoint, females had higher preference scores than males (p = 0.037). When we examined individual partner preference behaviors, we found that, at the group level, females and males huddled with their partner comparable amounts at the short-term timepoint (p = 0.45). Further, there was no consistent trend regarding which member of a pair huddled more with their partner; in ten pairs, the female huddled more and in six pairs, the male huddled more (Fig 1D). However, at the long-term timepoint, females consistently huddled more than their male partner (F > M in 15 of 16 pairs; p = 3.2×10^-5^). In the one pair that the male huddled more than the female, the difference in huddle times was negligible (less than 1% of the average huddle time for that pair). In contrast, female and male voles spent the same amount of time huddling with the novel at both timepoints (Fig 1E, short-term: p = 1.00, long-term: p = 0.54). Together, these data demonstrate that coordinated, intra-pair sex differences in affiliative behavior emerge as bonds strengthen and result from changes in female partner-directed huddle, but not male, behavior.

We next examined the time spent in partner and novel chambers when the test animal was not huddling (chamber time - huddle time), which serves as a proxy for investigative behavior. At the short-term timepoint, females spent more time investigating their partner than males did, but this difference was no longer evident at the long-term timepoint (Fig 1F, short-term: p = 0.028, long-term: p = 1.00). At both timepoints, males spent more time investigating the novel animal than females (Fig 1G, short-term: p = 0.0047, long-term: p = 6.8×10^-4^). While these behaviors differed by sex within pairs, there was no main effect of pairing time on within-sex behavior. Thus, these sex differences likely either reflect innate female/male differences or emerge extremely rapidly after pairing and do not reflect intra-pair coordination as there is no consistent relationship between female and male behavior. Finally, there were no within-pair sex differences or timepoint differences in the amount of time spent in the center chamber (Fig 1H) or in locomotion (Supplemental Table S1).

To examine our PPT data more holistically, we used principal component analysis (PCA) to aggregate and identify patterns within the following independent metrics: partner huddle time, novel huddle time, partner investigation, novel investigation, center chamber time, average distance to partner while in partner chamber, average distance to novel while in novel chamber, and total distance traveled. Using PCA to visualize the differences between sexes at the two timepoints, we found that at the short-term timepoint, female and male points largely overlap, and neither sex clusters together nor apart from the other sex (Fig 1I). However, by the long-term timepoint, females cluster together and apart from males, while males remain relatively dispersed (Fig 1J).

To determine which PPT metrics were driving each principal component, we performed a factor extraction, focusing on factors with a loading value > 0.3, indicating that 30% of the variance in that variable is explained by the principal component. At both timepoints, there is notable consistency in the specific behavioral factors that drive each principal component (Fig 1I, J). Specifically, PC1 is driven by partner and novel huddle time, novel investigation, and average within-chamber distance to the novel, with addition of partner investigation at the long-term timepoint. At both timepoints, PC2 is driven by partner huddle time, partner investigation, average within-chamber distance to partner, total distance traveled, and center chamber time. Finally, PC3 is driven by partner investigation, novel investigation, average within-chamber distance to novel, and center chamber time. At the long-term timepoint, PC3 is driven by partner investigation, total distance traveled, and center chamber time. Altogether, the first and second principal components broadly represent novel versus partner-directed behaviors, respectively, and this remains consistent over time.

Our analysis of our PPT data indicates that pair bonds develop through different behavioral changes in females and males and that behavioral sex differences emerge as bonds mature.

### Non-choice free interaction tests reflect partner preference and dyadic behavior

While PPT provides a valuable means to assess an individual’s behavior in the context of social choice, social interactions in the wild are not independently constrained to the actions of one individual. Thus, we also performed non-choice sequential free interaction tests, where we placed each animal, untethered, in a chamber with their partner or a novel (randomly ordered), allowing them to freely interact for 30 minutes. Free interaction tests were performed upon the animals’ initial introduction (baseline), and then again the day after short- and long-term PPTs (Fig 2A, B). In this free interaction test, huddling was qualitatively much less common than in the PPT, which may reflect the shorter duration of the test, limitations placed on social behavior due to tethering, and/or huddling as a form of consolation in prairie voles (Burkett *et al*. 2016). In addition, as there was no consistent way to parse the direction of interaction (e.g. male to female directed or vice versa), we scored total interaction time for each dyad (pair or each partner + novel).

**Figure 2:**
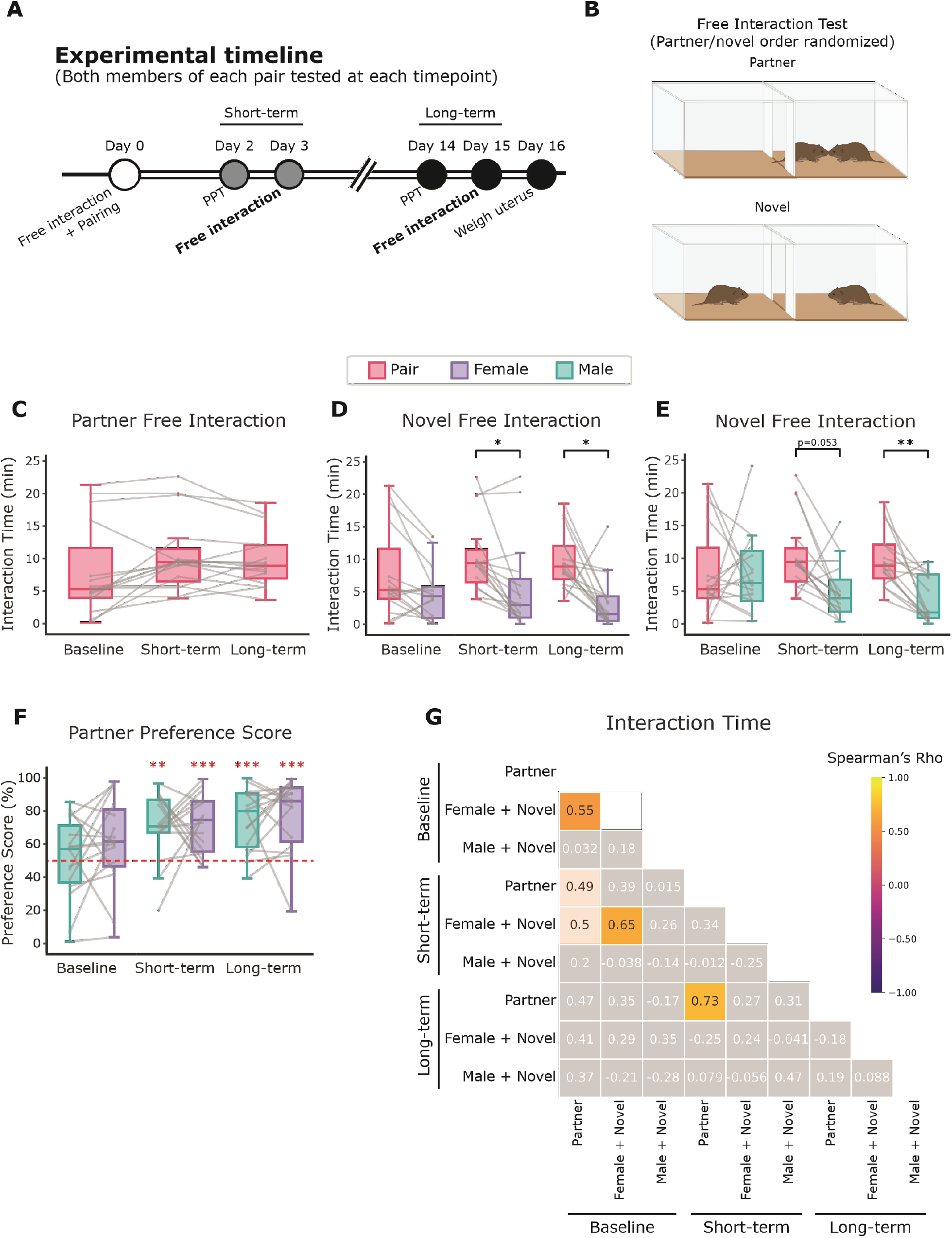
Non-choice free interaction tests as a measure of partner preference. **A.** Schematic of experimental timeline. Free interaction tests were conducted at short-term (3 days) and long-term (15 days) post-pairing. **B.** Diagram of free interaction tests. Animals were placed in an open chamber and allowed to freely interact with a partner or novel animal for 30 minutes. After an inter-test interval of at least 30 mins, the focal vole was tested with the other partner/novel (order randomized). **C.** Free interaction time between partners at baseline (day 0), short-term (day 3), and long-term (day 15) timepoints. No significant differences in pair interaction across timepoints. **D.** Female free interaction. Females interact more with their partner vs. a novel at the short- and long-term but not baseline timepoint (Baseline: p = 0.16, short-term: p = 0.025, long-term: p = 0.013). **E.** Male free interaction. Males interact slightly more with their partner at the short-term timepoint, which is more robust at the long-term timepoint (short-term: p = 0.053, long-term: p = 0.0047). **F.** Free interaction partner preference score calculated as Pair Interaction / (Pair Interaction + Novel Interaction) for each animal at each timepoint. Females and males show a significant partner preference at short- and long-term timepoints (females short-term: 1.9×10^-5^, females long-term: 1.3×10^-4^, males short-term: 0.0010, males long-term: 8.0×10^-5^). **G.** Correlation matrix of free interaction metrics between timepoints calculated using Spearman’s rho. Significant correlations are colored according to Rho value: baseline female + novel vs. baseline partner (Rho = 0.55, p = 0.027), baseline female + novel vs. short-term female + novel (Rho = 0.65, p = 0.0061), short-term partner vs. long-term partner (Rho = 0.73, p = 0.0013). Correlations with nominal significance (p < 0.06) are colored but opaque: baseline partner vs. short-term partner (Rho = 0.49, p = 0.052), baseline partner vs. short-term female + novel (Rho = 0.50, p = 0.050).

We found that at the group level, there are no differences in pair interaction time between timepoints (Fig 2C). However, while the majority of pairs (10 of 16) show modest increases in interaction between the baseline and long-term timepoints, six pairs decrease their interaction between these timepoints. Strikingly, four out of the six pairs that decrease are the same four pairs that exhibit notably higher levels of interaction than other pairs at the baseline timepoint. Despite this within-pair decrease, three of these pairs remain those with the highest interaction times at the long-term timepoint. Further, the percent change between the baseline and long-term timepoints is highly correlated with the amount of time spent interacting in the baseline test (Spearman’s Rho = −0.90, p = 1.7×10^-6^). Thus, although there is notable behavioral diversity between pairs, this demonstrates that within-pair behavior may change, but the pair’s behavior relative to other pairs remains consistent over time.

We next asked whether partner preference was evident in our free interaction paradigm by comparing the amount of interaction time with the partner and with the novel. Both males and females spent more time interacting with a partner than a novel animal at the short- and long-term timepoints (Fig 2D, E) indicating that sequential free interaction tests provide a valid indicator of partner preference. We further calculated a free interaction partner preference score (Pair Interaction / (Pair Interaction + Novel Interaction)) for each animal (Fig 2F). In this paradigm, both females and males show a partner preference by the short-term timepoint, which is maintained at the long-term timepoint. Compared to the PPT, this test did not reveal the same sex differences related to the strengthening of bonds over time, which may be due at least in part to the inability of this test to isolate behavior of one member of an interaction dyad.

We then asked whether there are any correlations between pair and novel free interactions to delineate which behavioral features correspond with bonding and which may reflect individual or sex-based differences in non-discriminate sociality. We used Spearman’s Rho to calculate correlation coefficients to avoid assumptions of linearity and account for order effects within the data, which is important for addressing behavioral consistency (e.g. do the pairs that spend the most time interacting at short term also do so at long term?). Only 3 of 36 potential correlations met a significance threshold of p < 0.05 as indicated by the colored boxes in Fig 2G. Correlations with p < 0.06 are colored but translucent. Specifically, we found that, at baseline, female interaction with their future partner or with the “novel” male was positively correlated (Rho = 0.55, p = 0.027), suggesting that some females may simply be more social than others, regardless of male interaction partner (Fig 2G). Notably, this was not true for males. Similarly, we found that female novel social interaction is positively correlated between baseline and short-term timepoints (Rho = 0.65, p = 0.0061), but neither baseline nor short-term is correlated with the long-term timepoint, indicating that this general sociality erodes as pair bonds mature (Fig 2G). We also found that partner social interaction is weakly correlated between baseline and short-term timepoints (Rho = 0.49, p = 0.052, Fig 2G), and, much more strongly, between short-term and long-term timepoints (Rho = 0.73, p = 0.0012, Fig 2G). This demonstrates enhanced intra-pair consistency over time with some pairs showing more interaction than others.

### Female, but not male, behavior correlates with pair behavior

We next determined the relationship between behaviors exhibited in the free interaction tests and PPTs using Spearman’s Rho. We calculated and included metrics that are likely to represent similar behavioral components across tests. Specifically, we reasoned that interaction in the free interaction test was likely most similar to the time an animal chose to interact with a tethered vole when it was near that animal. Thus, we calculated the huddle ratio – the percent time in the partner or novel chamber spent huddling (i.e. Huddle Time / Chamber Time). In addition, we calculated the within-pair Euclidean distance from the PCA of PPTs at each time point as a comprehensive indicator of within-pair similarity across multiple PPT metrics with a greater distance between partners representing more disparate behavior (Fig 1I, J, Fig 3A).

**Figure 3:**
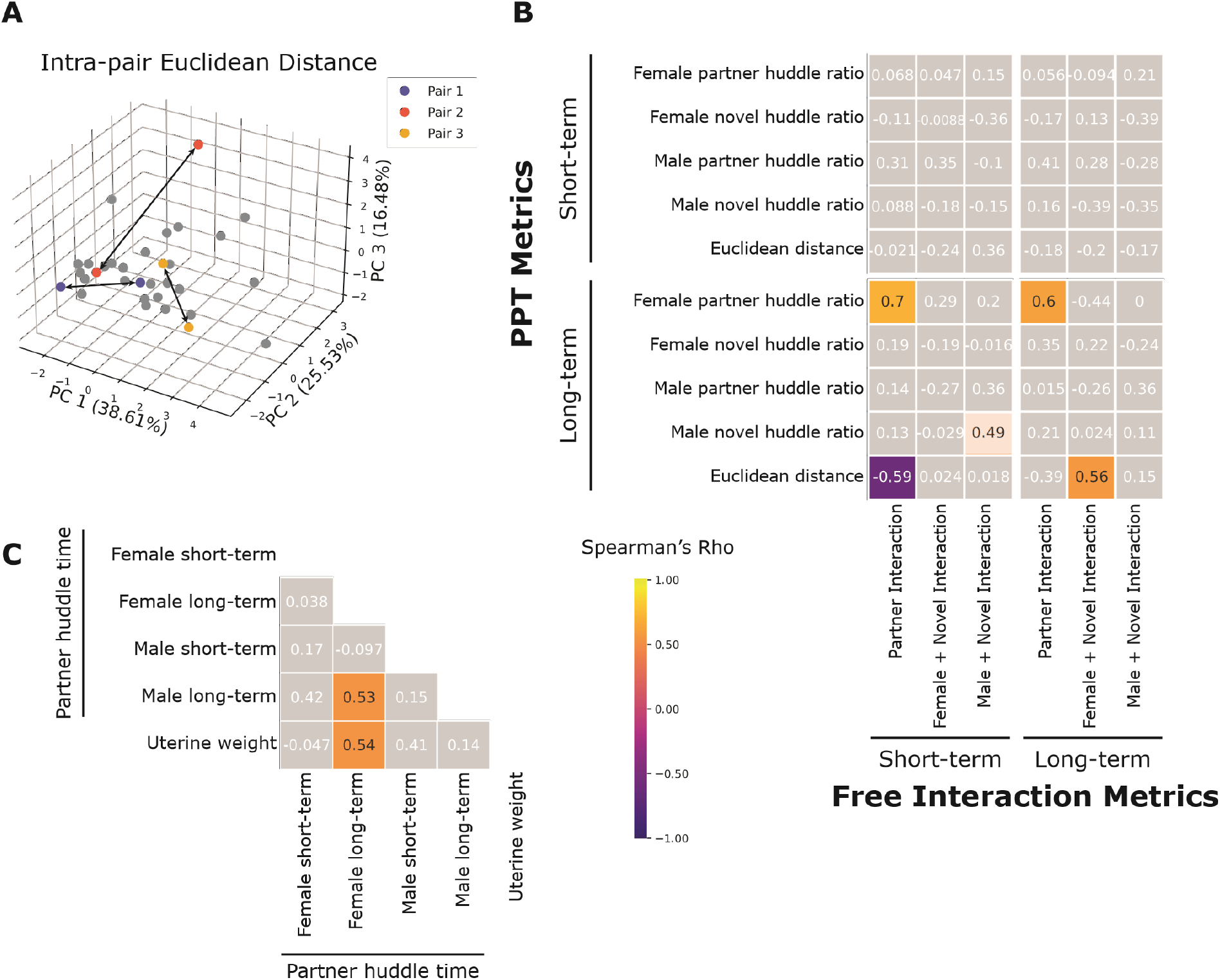
Correlations between PPT and free interaction test. **A.** Diagram of how Euclidean distance was calculated between partners within the same pairs from the PCAs in Fig 1I, J. **B.** Spearman’s Rho correlations between PPT and free interaction tests. Huddle ratio was calculated as Huddle Time / Chamber Time. Short-term PPT and free interaction test metrics did not correlate. The following metrics correlated significantly between the long-term PPT and short-term free interaction tests: female partner huddle ratio vs. partner free interaction (Rho = 0.70, p = 0.0027), PCA Euclidean distance vs. partner free interaction (Rho = −0.59, p = 0.015). The following almost reached significance: male novel huddle ratio vs. male + novel free interaction (Rho = 0.49, p = 0.055). Significantly correlated metrics between the long-term PPT and long-term free interaction tests include: female partner huddle ratio vs. partner free interaction (Rho = 0.60, p = 0.013), Euclidean distance vs. female + novel free interaction (Rho = 0.56, p = 0.024). **C.** Correlations between female and male PPT partner huddle times and uterine weight. The following correlations are significant: long-term female partner huddle vs. long-term male partner huddle (Rho = 0.53, p = 0.035), long-term female partner huddle vs. uterine weight (Rho = 0.54, p = 0.031).

We found that all PPT and free interaction metrics were uncorrelated at the short-term timepoint. However, correlations between these test metrics emerged by the long-term timepoint, suggesting a stabilization of a pair’s behavioral structure that emerges as a function of bond maturation. We found that at the long-term timepoint, pair interaction in the free interaction test is correlated with female partner huddle ratio, indicating that females that interacted more with their partner in the free interaction test also preferred to huddle when in proximity to their partner in the PPT. (Rho = 0.60, p = 0.013, Fig 3B. Additionally, female novel interaction in the free interaction test is positively correlated with the Euclidean distance between partners (Rho = 0.56, p = 0.023, Fig 3B). In other words, among pairs with lower intra-pair behavioral similarity in the PPT (a larger Euclidean distance), the female spends more time interacting with the novel in the free interaction test.

In addition to within-timepoint correlations, we also found that aspects of PPT behavior at the long-term timepoint correlated with a subset of metrics in the free interaction test at the short-term timepoint. This may suggest that even at the earlier bonding timepoints some behaviors are beginning to stabilize and are predictive of future behavior. The partner interaction time at the short-term timepoint was negatively correlated with the Euclidean distance between partners at the long-term timepoint (Rho = −0.59, p = 0.015, Fig 3B); the more time the pair spent together in the short-term free interaction test, the smaller the Euclidean distance between partners at the long-term timepoint. Partner interaction at the short-term timepoint was also correlated with female partner huddle ratio in the long-term PPT (Rho = 0.70, p = 0.0027, Fig 3B). Unlike females, male PPT partner behavior does not correlate with pair free interaction behavior. However, male free interaction with the novel at the short-term timepoint weakly correlates with male novel huddle time in the long-term PPT (Rho = 0.49, p = 0.055, Fig 3B). Taken together, our data suggest that female behavior can predict pair behavior specifically, while male behavior does not.

### Affiliative behavior as a function of pregnancy status

Finally, previous work suggests that pregnancy status can influence bond-related behaviors (Curtis 2010). Thus, we aimed to uncover any correlations between pregnancy status and behavior. The majority of pairs (15 of 16) became pregnant during the two weeks of pairing. At 16 days post-pairing, females were sacrificed, uteri were weighed, and embryos were weighed and counted. We found that uterine weight was positively correlated with raw female partner huddle time (Rho = 0.54, p = 0.031), but not male partner huddle time (Rho = 0.14, p = 0.61), at the long-term PPT (Fig 3C). While raw male partner huddle time was not correlated with uterine weight, male partner huddle time was correlated with female partner huddle time at the long-term timepoint (Rho = 0.53, p = 0.035). Interestingly, our one pair that did not become pregnant over the course of our experiment is the same pair that showed no sex difference in partner huddle time in the long-term PPT. Together, this suggests that pregnancy status may alter female behavior which, in turn, may drive changes in male behavior.

### Females display greater partner-directed motivation than males

To determine if sex differences observed in PPT and free interaction behavior may be partially explained by differences in selective social motivation, we trained 6 female and 6 male prairie voles to press for social access. To confirm that voles had bonded with their sterilized mates, we performed a three hour PPT. As previously observed (Fig 1C), male voles displayed more variability in their partner preference scores, with 2 of 6 males displaying a novel preference, and 3 of 6 displaying scores greater than 80%. In contrast, all females displayed preference scores greater than 80% (Fig 4C). While not statistically significant, females displayed greater partner huddle times than males, consistent with earlier observations (Fig 4D, 1D, long-term).

**Figure 4:**
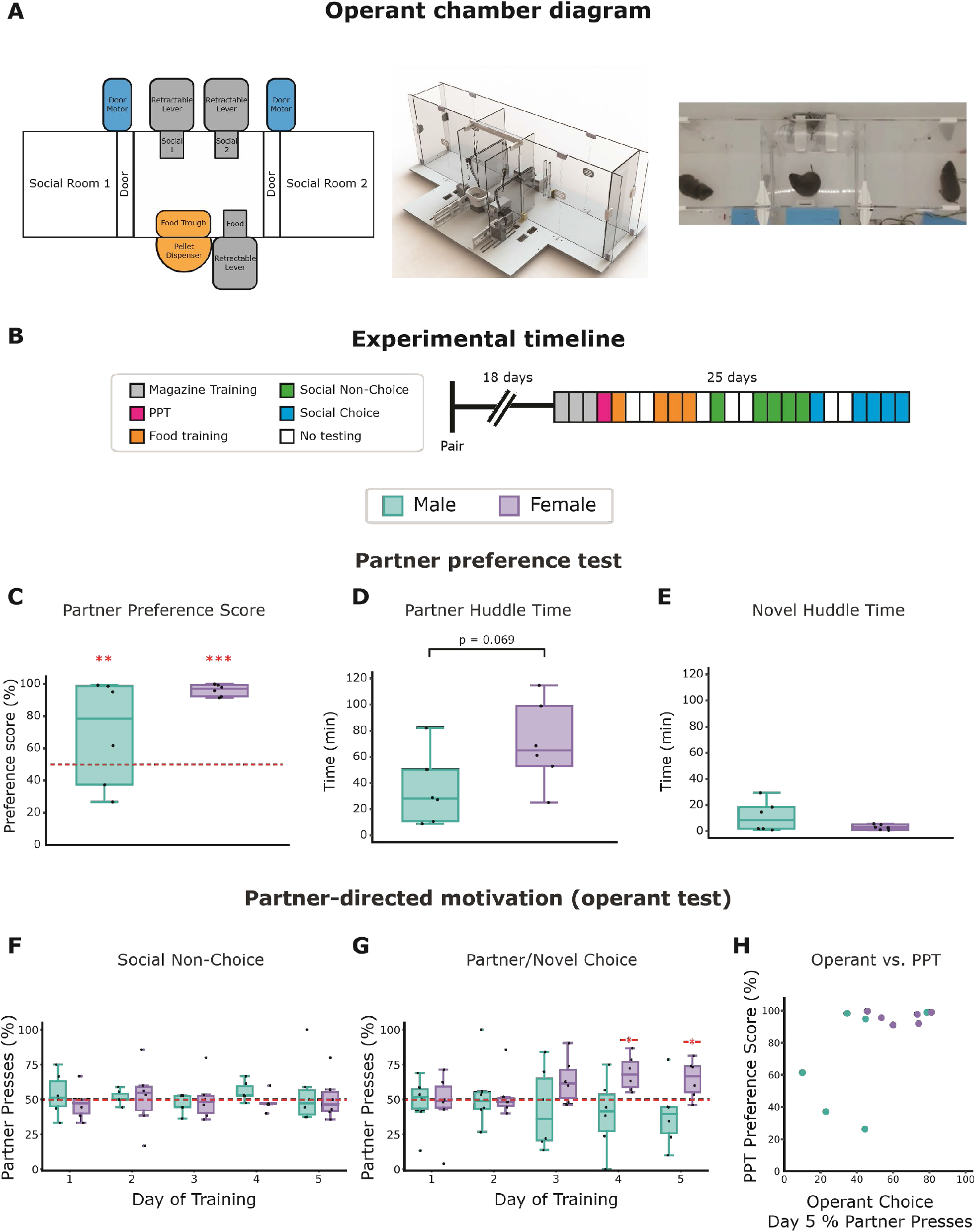
Operant paradigm for assessing partner-directed motivation. **A.** Social choice operant apparatus. Left: schematic of relevant components for lever delivery and access to food or social reward. Middle: 3-dimensional diagram of apparatus designed in Solidworks and visualized in Photoview 360. Right: Top down screenshot of social operant task. **B.** Experimental timeline. Voles learned to associate relevant cues with food pellet delivery during magazine training (gray boxes), and then underwent training in which they received a food pellet faster if they pressed the lever (orange boxes). This was repeated for access to a social reward (partner or novel alternated in 5 trial bins; green boxes). Finally, social choice was assessed via an exclusive choice task in which both levers were presented and the test animal could receive access to either the partner or novel animal during each trial (blue boxes). **C.** Partner preference scores for test conducted 3 weeks post-pairing (pink box in B), calculated for each animal as Partner Huddle Time / (Partner Huddle Time + Novel Huddle Time) x 100%. Red asterisks denote significant difference from the null hypothesis of no partner preference (50%) using a one-sample t-test (females: p = 1.7×10^-8^, males: p = 0.0036). Females show a non-significant increase in preference score compared with males (t-test, p = 0.080) **D.** Females huddled more with their partner than males (t-test, p = 0.069). **E.** No sex differences were observed in novel huddle duration. **F.** When one lever is presented at a time, male and females will press equally for access to the partner or the novel. **G.** In a choice paradigm, a preference for partner access emerges by testing day 4 for the females but not the males. Red asterisks denote significant difference from the null hypothesis of no preference (50%) using a one-sample t-test (females: Day 4 p = 0.014, Day 5 p = 0.047). **H.** Scatterplot showing a general separation in female and male behavior based on partner preference score and percent partner lever presses in the operant choice paradigm.

Prairie voles rapidly learned to press for food pellets in our training paradigm, with most animals pressing on more than 50% of trials after 3 days of magazine training, and 4 days of food training (males: 4 of 6, females 5 of 6). Additionally, across training days both males and females begin to press the lever more quickly after the lever is extended at the start of a round (Fig S1A, main effect of day, p = 0.0021). With time, they increase the number of rounds in which they successfully lever-pressed (Fig S1B, main effect of day, p = 0.017), both indicating that the animals learned the task.

Following food training, voles were trained to press for social access via a similar paradigm. The partner lever was extended 5 trials in a row followed by opening of the partner door 30 seconds later, with the option to open the door sooner if the test vole pressed the lever. The door remained open for the rest of the trial, so if a vole pressed immediately upon the start of the trial, it received up to 90 seconds of social access, while failure to press still resulted in 60 seconds of social access. The animal was then presented with the novel lever 5 trials in a row, and this pattern was repeated once for a total of 20 trials. Animals pressed more frequently after the first day (Fig S1C, main effect of day, p = 0.0091), indicating they learned the task consistent with prior reports (Matthews *et al*. 2013; Beery *et al*. 2021). Interestingly, even though most animals display a preference in the PPT, we did not observe a preference in pressing when presented with only one lever at a time (Fig 4F).

In order to more directly ask if prairie voles displayed differential motivation to access their partner versus a novel vole, we then presented trained animals with both social levers simultaneously across 30 trials. On any trial, the choice between partner and novel was mutually exclusive. Females reliably demonstrated that they had learned this task by the 5th day of testing, demonstrated by correctly orienting towards the selected door prior to the door opening. (Fig S1E, F, one sample t-test vs. 0.5, p = 3.1×10^-4^), Males, as a group, were not better than chance (p = 0.074), although this was primarily driven by one individual (Fig S1F). When removed, the remaining 5 males also correctly oriented more frequently than chance (p = 0.0065 excluding vole 4139).

We found that female voles changed their pressing behavior over time, displaying a statistically significant preference to lever-press for their partner on days 4 and 5 (one sample t-test vs. 50%, Day 4: p = 0.014, Day 5: p = 0.047). Conversely, male vole lever-pressing did not indicate a partner or a novel preference, and on day 5 only 1 of 6 males pressed more for their partner (Fig 4G, Day 5: p = 0.45). This observation in female voles contrasts with the lack of preference observed in the non-choice training phase (Fig 4F). As selective social motivation likely impacts preference in the PPT, we compared PPT preference scores to operant choice pressing preference (Fig S1D) and found that males tended to score lower than females for both metrics.

Finally, we asked if males and females showed different behaviors after pressing to gain social access. Although we did not observe any statistically significant differences between males and females, their post-pressing behavior trends match those observed in pairs in the PPT (Fig 1), with a trend for females to huddle more with their partners (p = 0.102) and a trend for males to spend more non-social time in the novel chamber (Fig S1G, p = 0.088). Males and females displayed similar levels of aggression, with greater aggression directed towards the novel than the partner, although this did not reach threshold for significance (Fig S1H, main effect of interaction partner, p = 0.134).

## Discussion

The vast majority of studies examining sex differences in prairie voles do so in the context of parenting behavior. While important, this leaves us with a relative lack of understanding of behavioral sex differences that may be critical to forming and maintaining a bond pre-parenting. Critical work performed over two decades ago demonstrated that partner preferences develop more rapidly post-pairing in females than males (DeVries & Carter 1999). However, whether sex differences are seen within individual pairs, and whether it is consequential to facilitate coordinated behavior, has remained unexplored.

### Choice and non-choice social tests reveal female-driven behavioral coordination

Here, we provide the first intra-pair comparisons of affiliative pair bonding behavior in prairie voles. We tested both members of female/male pairs in PPTs and in free interaction tests at short- and long-term pairing timepoints. We first found that pair bonds mature over time via different mechanisms in females and males, with females increasing their partner affiliative behavior, and males decreasing their novel affiliative behavior. Further, we found that sex differences and coordinated behavior between partners emerge as a pair bond matures, an observation bolstered by female/male differences in clustering in our principal component analysis. Our data point to the importance of testing pair bond behaviors over the life of a bond, as bond formation and maturation represent different stages underwritten by distinct behavioral mechanisms.

We next tested prairie vole pairs in sequential free interaction tests, a non-choice paradigm in which the experimental animal has the option to either interact with their untethered partner or a novel in the absence of the other. Despite being a non-choice test, the free interaction test recapitulated results from the PPT, with animals choosing to spend more time interacting with their partner than a novel. Unlike the PPT, the free interaction test uniquely allowed us to test dyadic pair bonding behavior — behavior resulting from the actions of both partners at the same time — rather than isolating the partner-directed behavior of one animal as in the PPT.

When we compared individual partner-directed behavior in the PPT and dyadic pair behavior during free interaction, we found that female partner-directed behavior correlates with pair behavior, while male behavior does not. Once in proximity to their partner in the PPT (e.g. in the partner chamber), the experimental animal has the choice to either interact with their partner or not, which is analogous to the choice to interact with a non-tethered animal in the free interaction test. Thus, we calculated the partner huddle ratio (Partner Huddle Time / Partner Chamber Time) from the PPT and found that female partner huddle ratio at the long-term timepoint is correlated with pair free-interaction at both the short- and long-term timepoints. To further compare our two tests, we used PCA to reduce the dimensions of our PPT data and then calculated the Euclidean distance between partners within the same pair. Interestingly, we found that the Euclidean distance between partners at the long-term timepoint was inversely correlated with partner interaction at the short-term timepoint. As a larger Euclidean distance between partners represents more disparate behavior in the PPT, our data indicate that less pair interaction at the short-term timepoint predicts more dissimilar behavior at the long-term timepoint. In addition, Euclidean pair distance is positively correlated with female::novel free interaction at the long-term timepoint indicating that in the pairs with more dissimilar behavior in the PPT, the female spends more time interacting with the novel in the free interaction. Across the PPT to free interaction comparisons, female behavior correlated with pair behavior while male behavior did not. Together, this suggests that female behavior is a primary driver of pair behavior and therefore behavioral coordination.

### Emergent coordinated sex differences serve a function other than mate choice

Sex differences are thought to exist primarily for two intertwined purposes: mate choice and reproduction (Darwin 1871; Andersson & Iwasa 1996; Price 1998). Compared to promiscuous species, monogamous species typically exhibit fewer sex differences (Kleiman 1977), and in our experiment, pairs were randomly pre-assigned. Together, this strengthens the argument that the sex differences that emerge as pair bonds form and mature serve a different function than those that exist to drive mate choice and sexual selection.

Instead, emerging sex differences in pair bonding behaviors may help prime pairs to co-parent. Notably, our observed sex differences are coordinated within pairs. For instance, intra-pair affiliation (partner huddle) is not simply higher in females, it is correlated within pairs. Supporting a role for this behavioral coordination in future parenting, female huddle is correlated with uterine weight, and as such, pregnancy status may be driving correlated male-female behavior by acting as a set-point for female partner huddle which the male uses to calibrate his behavior. Interestingly, the one pair that did not become pregnant over the course of our experiment was also the one pair with inconsistent female/male behavior; the female did not show greater partner huddle time than her male partner in the long-term PPT. While it is unclear whether the lack of pregnancy drives the lack of coordinated behavior or vice versa, it does support a broad role for behavioral coordination in facilitating reproduction. Notably, similar mechanisms in which hormones drive female behavior which, in turn, changes male behavior have been observed in pair bonding bird species (Martinez-Vargas & Erickson 1973).

### Sex differences in partner-directed motivation broadly reflect differences in partner affiliation

Using a separate cohort of animals, we dually employed social operant testing and PPTs. Replicating our previous experiment, there were notable sex differences in partner huddle time, with females huddling more with their partners than males. When provided with a fixed ratio of one lever press per social reward, in which voles pressed separately for either a partner or a novel in separate trials, females and males pressed equally for access to their partner and a novel vole in this testing paradigm. Although this differs from findings by Beery *et al*. (2021) showing that female prairie voles work harder to access familiar males in a non-choice operant task, there are two important differences. Beery et al. employed a progressive ratio in which voles had to increase pressing across trials to get the same social access, and if they failed to press, they did not gain access. In contrast, our paradigm required substantially less effort and voles gained access to the social reward even if they did not press. Thus, it is likely that sex differences are only evident under increasing task demands or in a choice context. Accordingly, when given a choice of pressing to gain access to their partner or a novel vole, females showed biased pressing to access their partner more frequently than accessing the novel. These sex differences in partner-directed motivation are broadly reflected in the sex differences in PPT partner-directed affiliation (although these two metrics are not strongly correlated at the individual level). Given that females huddle with their partner more than males across both PPT experiments and in the operant paradigm, sex differences in partner-directed motivation may partially be responsible for increased affiliative behavior and, by extension, pair behavior. The present findings in the partner/novel operant choice test mirror those of similar tests conducted by Vahaba *et al*. (also submitted to this issue). Of particular note, the results reported by Vahaba et al. and in this manuscript are consistent despite differences in testing apparatus, training paradigms, food restriction, colony origins, altitude, and other factors, indicating that sex differences in partner-directed effort are highly robust.

Notably, our operant experiment was designed to assess sex differences but not within the context of a pair. This was largely due to constraints related to the daily training and testing required for this experiment. Further, the longitudinal nature of this experiment required sterilization of the untested partner. This experiment therefore indicates that pregnancy itself is not required for emergence of behavioral sex differences, although whether it is required for the intra-pair coordination of these behaviors remains unknown.

### Limitations and future directions

One limitation of the current work stems from the relatively sparse schedule of testing (3 timepoints), which occurred entirely before females were due to give birth. Future work is needed to resolve the time course of behavioral changes, pair-based variability, and the factors that may drive this variation. Likewise, although coordinated behavior is evident in affiliative and parenting behavior, it remains unknown whether coordinated behavior in these two tasks is related. Manipulations of pregnancy status and/or pup presence may provide fruitful insight into this question.

Our data demonstrate that prairie voles exhibit behavioral sex differences that contribute to reliable patterns of intra-pair behavior. These sex differences emerge as a bond matures and correlate with pregnancy. This emergent coordination may serve to facilitate future biparental care, or may simply be a result of cumulating social experience between two individuals. Together, this work and future work can uncover how coordinated behavior arises between bonded partners and its role in promoting success of a species.

## Materials and Methods

### Animals

Adult prairie voles were bred in-house in a colony descended from wild animals collected in Illinois. After weaning at 21 days, animals were housed in same-sex groups of 2-4 animals in standard static rodent cages (17.5 l. x 9.0 w. x 6.0 in. h.) with ad-libitum access to water and rabbit chow (5326-3 by PMI Lab Diet). Diet was supplemented with sunflower seeds and alfalfa cubes, and cotton nestlets and plastic houses were given for enrichment. All voles were between 58 and 90 days of age at the start of the experiment. Beginning on day one, female/male pairs were co-housed in smaller static rodent cages (11.0 l. x 6.5 w. x 5.0 in. h.) with ad-libitum access to water and rodent chow, as well as cotton nestlets and houses for enrichment. Animals were kept at 23-26C with a 14:10 light:dark cycle. All procedures were performed during the light phase and approved by the University of Colorado Institutional Animal Care and Use Committee.

### Timeline

Experimental timeline shown in Figure 1A was carried out for 16 female/male vole pairs. Briefly, baseline tests (day 0) consisted of two free interaction tests: one with the animal they would subsequently be paired with, and one with an animal they would not be paired with (“novel”). After all 30-minute free interaction tests were complete, animals were co-housed with their randomly pre-selected partner for the duration of the experiment. At 2 days post-pairing, we performed short-term tests consisting of partner preference tests (Williams 1992) (PPTs) performed sequentially for both animals within each pair followed by free interaction tests one day later (day 3). The pairs continued to cohabitate and were tested at a long-term timepoint via sequential PPTs on day 14 and free interaction tests on day 15. Each animal was tested with a different novel animal for each test, ensuring that the animals never saw the same novel animal twice. At 16 days post-pairing, animals were sacrificed to weigh the uterus and count embryos. Across all tests, test order for female and male was randomized, and for free interaction tests, the order of partner or novel presentation was randomized to account for potential order effects.

### Free interaction

Free interaction tests were performed in clear rectangular plexiglass arenas 50.7 cm long, 20.0 cm wide, and 30.0 cm tall. For each test, experimental animals were paired either with their partner or a novel other-sex animal, order randomized. All animals had an inter-trial interval of 30-90 minutes. Animals were individually placed on opposite sides of the chamber separated by an opaque divider. At the start of the test, the divider was removed and both animals were allowed to freely move about the chamber for 30 minutes. Overhead cameras (Logitech C925e webcam) were used to record four free interaction tests simultaneously.

Periods of social interaction between the two animals were scored post-hoc using TopScan High-Throughput software v3.0 (Cleversys Inc). We adapted and optimized scoring methods from (Ahern *et al*. 2009) and defined social contact by setting the “joint motion” parameter to < 5. To confirm the accuracy of the TopScan software, two pairs were hand-scored using BORIS (Friard & Gamba 2016) for the following behaviors: interacting, affiliative behavior, neutral behavior, and aggressive behavior. Compared to the amount of interacting time scored by hand, the TopScan-scored interacting time differed by less than 6% in both videos.

### Partner Preference Test

Partner preference tests were performed as described in Scribner et al. 2020 (Scribner *et al*. 2020). Briefly, both partner and novel animals were tethered to the end walls of three-chamber plexiglass arenas (76.0 cm long, 20.0 cm wide, and 30.0 cm tall). Tethers consisted of an eye bolt attached to a chain of fishing swivels that slid into the arena wall. Animals were briefly anesthetized with isoflurane and attached to the tether using a zip tie around the animal’s neck. Two pellets of rabbit chow were given to each tethered animal and water bottles were secured to the wall within their access while tethered. After tethering the partner and novel animals, experimental animals were placed in the center chamber of the arena. At the start of the test, the opaque dividers between the chambers were removed, allowing the experimental animal to move freely about the arena for three hours. Overhead cameras (Panasonic WVCP304) were used to video record eight tests simultaneously.

The movement of all three animals in each test was scored using TopScan software using the parameters from Ahern *et al*. 2009. Behavior was analyzed using a Python script developed in-house (https://github.com/donaldsonlab/Cleversys_scripts) to calculate the following metrics: time spent in partner/novel chamber, time spent huddling with partner/novel, average distance to partner/novel while in the respective chamber, latency to huddle with partner/novel, and total locomotion. The partner preference score was calculated as (Partner Huddle Time / (Partner Huddle Time + Novel Huddle Time)) x 100%.

### Weighing the uterus/counting embryos

Following the final free interaction test, animals were sacrificed to weigh the uterus and to measure embryo head-to-rump length. Animals were euthanized using CO_2_ and decapitation. Uteri were then dissected out from the females and weighed. From each uterus, embryos were counted and one embryo was removed to measure head-to-rump length.

### Statistics

Data were analyzed using the SciPy Stats package (Virtanen *et al*. 2020) (version 1.7.0) and Pingouin package (Vallat 2018) (version 0.3.12) in Python (version 3.8.10). Details of all statistical tests can be found in Supplemental Table S1. To determine the statistical significance of the partner preference score (i.e. whether a partner preference was formed), we used a one-sample t-test comparing to a value of 50% (no preference for partner/novel). To assess the intra-pair effects of sex and the within pair effects of time, we used repeated measures ANOVA (RM-ANOVA). Because the females and males are intrinsically paired, and within a pair, female and male behavior are not independent, we used paired t-tests with Bonferroni correction for our post hoc comparisons. For analysis of the PPT partner preference scores, we used Wilcoxon rank sum tests for our post hoc comparisons because the scores are not normally distributed. For our correlation analyses, we calculated all of our correlations using Spearman’s Rho to avoid assumptions of linearity and account for order effects, neither of which are possible using the more traditional Pearson’s R. Throughout the paper, * indicates p < 0.05, ** indicates p < 0.01, and *** indicates p < 0.001. In the correlation matrices, R values with associated p values < 0.05 are colored. Nominally significant correlations (p < 0.06) are colored lighter and slightly translucent.

### Open Source Custom Operant Chamber

Operant chambers contained 3 chambers separated by 2 motorized doors, 3 separate retractable levers, and one motorized pellet dispenser and trough (Fig 4A). Chambers were constructed from a mix of laser cut acrylic and 3D printed ABS plastic. A bill of materials and chamber designs can be accessed at https://github.com/donaldsonlab/Operant-Cage/tree/main/V2.

The box was controlled via custom scripts and code (https://github.com/dprotter/RPi_Operant) run on Raspberry Pi computers (Raspberry Pi Foundation). Servos were controlled via an Adafruit HAT (Adafruit 2327). Each chamber was controlled by a corresponding Raspberry Pi. Food rewards were 20 mg pellets (Dustless Precision Pellets Rodent Grain-Based Diet; VWR 89067-546) delivered to a trough. Pellet dispensal and retrieval was detected by an IR beam break in the trough. Tones were generated via PWM on the Raspberry Pi (pigpio), and played through an amplified speaker (Adafruit 3885).

### Operant Timeline

Animals (n = 12, 6M, 6F) were trained using in-house constructed operant chambers to perform a social selection operant task. Partners for test animals were sterilized (tubally ligated or vasectomized as described in Donaldson *et al*. 2009 and Harbert *et al*. 2020) at least 2 weeks prior to pairing. Test animals were paired and cohabitated for 18 days before the start of operant training. Animals underwent 3 days of magazine training, 1 day of partner preference test, 4 days of food training, 5 days of social training, and 5 days of social choice testing. Animals were not trained or tested on weekends. Novel stimulus animals were rotated to minimize potential familiarity. All sterilized partners were used as novel stimuli, along with 5 additional unpaired, intact males and 3 unpaired, intact females.

### Magazine Training

Animals underwent 15 rounds in which a tone was played to indicate the start of the round (5,000 Hz, 1s). The food lever was then extended for 2 seconds, a pellet cue (2500 Hz, 1s) was played, and a pellet was delivered to the trough. The lever was retracted 2 seconds later. Lever presses immediately triggered the pellet reward, with a maximum of 1 reward per round. Total round time was 90s.

### Food Training

Animals underwent 15 rounds in which a tone was played to indicate the start of the round (5,000 Hz, 1s). The food lever was then extended for 30 seconds. After 30s, the lever was retracted, a pellet cue (2,500 Hz, 1s) was played, and a pellet delivered to a trough. If the animal pressed the lever the lever was retracted, pellet cue was played, and a pellet was immediately dispensed. In order to provide a window to observe anticipatory behavior, animals experienced a delay between lever pressing and reward as follows: [day 1: no delay, day 2: no delay, day 3: 1 s, day 4: 1 s]. Total round time was 90s.

### Social Training

Animals underwent 20 rounds of social training, in alternating sets of 5 trials for each door, starting with the partner door (5 partner, 5 novel, 5 partner, 5 novel). The location of the partner and novel animals was kept the same across days. Stimulus animals were tethered to the walls farthest from the doors in a similar fashion to the PPT. On each trial, a tone was played to indicate the start of the round (5,000 Hz, 1s). The corresponding social lever was extended for 30s. After 30s, the lever was retracted, a door-opening cue was played (10,000 Hz, 1s) and the corresponding door opened. If the lever was pressed, the door-opening cue was played and the door opened immediately. At the end of the trial a door close tone was played (7,000 Hz, 1s) and the door was closed. Total round time was 110s, with 20s allocated for researchers to return the test animal to the central chamber in between trials, if necessary. Therefore, all animals always received a minimum of 60s of partner or novel access on a trial, but animals that pressed more quickly received longer access. Delays between pressing and door opening were as follows: [day 1: no delay, day 2: no delay, day 3: 1s, day 4: 1s, day 5: 2s].

### Social Choice Testing

Animals underwent 30 rounds of social testing. The location of the partner and novel animals was kept the same as in social training. On each trial, a tone was played to indicate the start of the round (5,000 Hz, 1s). Both social levers were extended for 30s. If a lever was pressed, both levers were retracted, the door-opening cue was played (10,000 Hz, 1s) and the corresponding door opened after a 1s delay. If no lever was pressed within 30s, both levers were retracted. At the end of the trial a door close tone was played (7,000 Hz, 1s) and the door was closed. Total round time was 110s, with 20s allocated for researchers to return the test animal to the central chamber in between trials, if necessary. Therefore, animals that pressed more quickly received more social access, with a minimum of 60s social access. Only one lever-press was allowed per trial, making social access mutually exclusive during each trial.

### Data Sharing

Data that support the findings of this study will be made available via Figshare at the time of publication, and this section will be updated with a reference number. Complete statistical results, including effect sizes are reported in the supplementary Statistics Table. Code and operant apparatus design files are available via the GitHub links provided in the methods section.

## Acknowledgments

We thank Cayla Jo Paulson and Jessica Abazaris of the animal care staff at the University of Colorado Boulder. The majority of fabrication for custom operant apparatuses was conducted at the Integrated Teaching and Learning Laboratory at the University of Colorado Boulder. We thank the rest of the Donaldson lab for their feedback and support and the voles for their sacrifice. This work was supported by NIH award DP2OD026143, NSF IOS-2045348, NSF IOS-1827790 and funds from the Whitehall Foundation and the Dana Foundation (to Z.R.D.), NIH R15MH113085 (to A.K.B), and NIH T32 GM008759-17/18 (to L.E.B.).

## Author Contributions and Notes

L.E.B., D.S.W.P., A.K.B., and Z.R.D. designed research. L.E.B., D.S.W.P., A.C.F., M.U.P., and I.O.E. performed research. D.S.W.P, G.D.C., and R.T.C. designed and engineered operant chambers. L.E.B. and D.S.W.P. analyzed data. L.E.B., D.S.W.P., and Z.R.D. wrote the paper. The authors declare no conflicts of interest.

## Notes

### Competing Interest Statement

The authors have declared no competing interest.

