## Supplemental Figure 1 for "Emergent intra-pair sex differences and behavioral coordination in pair bonded prairie voles"

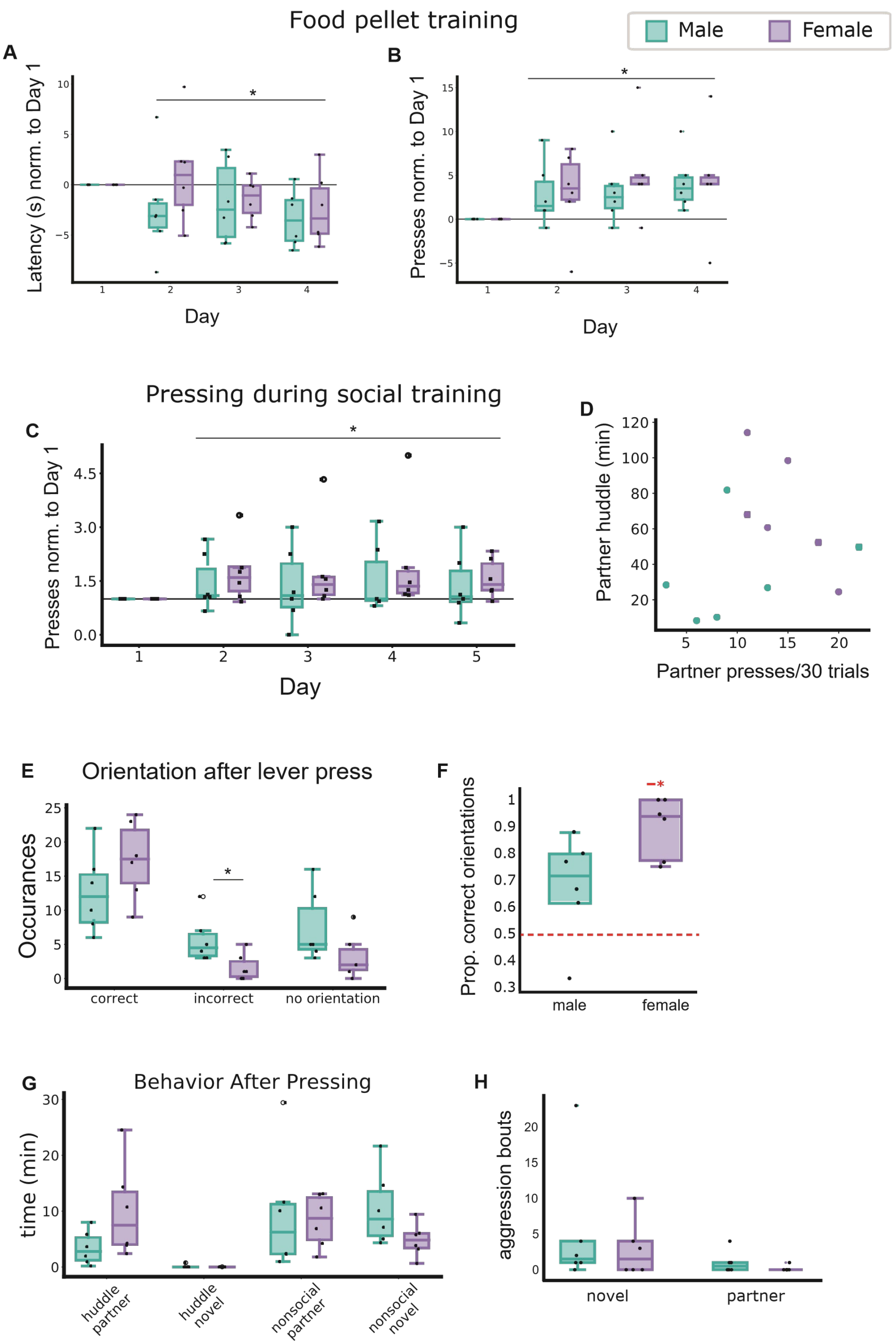

Food pellet training
Male
Female
A
Latency (s) norm. to Day 1
Day
B
Presses norm. to Day 1
Day
Pressing during social training
C
Presses norm. to Day 1
Day
D
Partner huddle (min)
Partner presses/30 trials
E
Orientation after lever press
Occurances
correct
incorrect
no orientation
F
Prop. correct orientations
male
female
G
Behavior After Pressing
time (min)
huddle partner
huddle novel
nonsocial partner
nonsocial novel
H
aggression bouts
novel
partner

**Supplementary figure 1. Additional behavioral metrics during operant testing. A, B.** Voles are distributed a food pellet 60 seconds after lever and cue presentation but can receive it faster by pressing the lever. Latency to press the lever decreases (A) and the lever is pressed in more trails across training days (B). **C.** Change in pressing relative to the first day of social training. **D.** Males and females separate out when examining the relationship between partner huddle in the PPT (Fig 4D) and number of times they pressed for their partner in the choice test on the last day of testing. **E, F.** We used orientation after pressing as a proxy for whether the animal had formed a reliable lever/door association. Data shown is from the last day of social training (E). Males and females oriented towards the correct door more often than they oriented towards the incorrect door, although this was only significant for females ($p = 0.00031$). For males, the lack of significance was due to a single animal who also reliably performed poorly in other task metrics ($p = 0.074$) (F). **G.** Post-pressing behavior from the last day of the social choice test. Females huddled more with their partner than males did after pressing for access. There was virtually no huddling with the novel vole. There were no sex differences in the time spent in the partner chamber but not actively engaged with the tethered partner, but males spent more time than females in the novel chamber but without physical interaction with the tethered vole, reminiscent of investigative behavior in the PPT (Fig 1G). **G.** Number of aggression bouts did not differ for males and females.
